# Tonic dopamine, uncertainty and basal ganglia action selection

**DOI:** 10.1101/2020.11.10.376608

**Authors:** Tom Gilbertson, Douglas Steele

**Affiliations:** Department of Neurology, Level 6, South Block, Ninewells Hospital & Medical School, Dundee, DD2 4BF, UK; Division of Imaging Science and Technology, Medical School, University of Dundee, DD2 4BF, UK

## Abstract

To make optimal decisions in uncertain circumstances flexible adaption of behaviour is required; exploring alternatives when the best choice is unknown, exploiting what is known when that is best. Using a detailed computational model of the basal ganglia, we propose that switches between exploratory and exploitative decisions can be mediated by the interaction between tonic dopamine and cortical input to the basal ganglia. We show that a biologically detailed action selection circuit model of the basal ganglia, endowed with dopamine dependant striatal plasticity, can optimally solve the explore-exploit problem, estimating the true underlying state of a noisy Gaussian diffusion process. Critical to the model’s performance was a fluctuating level of tonic dopamine which increased under conditions of uncertainty. With an optimal range of tonic dopamine, explore-exploit decision making was mediated by the effects of tonic dopamine on the precision of the model action selection mechanism. Under conditions of uncertain reward pay-out, the model’s reduced selectivity allowed disinhibition of multiple alternative actions to be explored at random. Conversely, when uncertainly about reward pay-out was low, enhanced selectivity of the action selection circuit was enhanced, facilitating exploitation of the high value choice. When integrated with phasic dopamine dependant influences on cortico-striatal plasticity, the model’s performance was at the level of the Kalman filter which provides an optimal solution for the task. Our model provides an integrative account of the relationship between phasic and tonic dopamine and the action selection function of the basal ganglia and supports the idea that this subcortical neural circuit may have evolved to facilitate decision making in non-stationary reward environments, allowing a number of experimental predictions with relevance to abnormal decision making in neuropsychiatric and neurological disease.

## Introduction

To make optimal decisions in uncertain circumstances, flexible adaption of behaviour is crucial; exploring alternatives when the best choice is unclear, exploiting what we know when we think that is best. Critical for optimally solving the “explore-exploit” dilemma is a decision making strategy which strikes a balance between these two approaches. A Kalman filter is optimal under certain conditions (Kalman, 1960) and evidence for human motor control (Orban de Xivry *et al*., 2013), perception and imagery (Grush, 2004) conforming to Kalman filter predictions has long been noted. However, the neural implementation of human optimal decision-making remains unclear.

Experimental studies have focused on the role of pre-frontal cortical circuits which differentially meditate explorative or exploitative decisions (Daw *et al*., 2006; Chakroun *et al*., 2020). Recent data suggest that subcortical regions, including the striatum, encode both exploratory and exploitative choices in their single neuron activity (Costa *et al*., 2019). Consistent with this finding is the recognised role for striatal dopamine in determining the explore-exploit trade-off (Zhuang *et al*., 2001; Frank *et al*., 2009a; Beeler *et al*., 2010; Costa *et al*., 2014). However, relatively little attention has been given to the question of whether the basal ganglia circuit, in relative isolation, can solve the explore-exploit problem. From a theoretical perspective, the basal ganglia circuitry, through its action selection function (Gurney *et al*., 2001) is ideally suited to provide both a flexible and precise solution (Humphries *et al*., 2012).

A widely used paradigm to investigate explore-exploit behaviour experimentally is the multi-armed “restless” bandit task (Gittins & Jones, 1979; Daw *et al*., 2006; Speekenbrink & Konstantinidis, 2015). In this paradigm, the reward pay-out for each of the choices, fluctuates on a trial-by-trial basis in accordance with a Gaussian diffusion process. To perform optimally, the participant under conditions of both high volatility and choice uncertainty, has to identify and track the highest value choice.

Here we test the idea that the decision to explore or exploit is governed in the basal ganglia by the *interaction* between tonic dopamine and cortical on selectivity of the basal ganglia’s action selection mechanism (Humphries *et al*., 2012; Suryanarayana *et al*., 2019). In accordance with experimental predictions (Fiorillo *et al*., 2003; St Onge *et al*., 2012) we show that tonic dopamine levels track the uncertainty of the reward pay-out. Under increased uncertainty, higher levels of tonic dopamine led to increased random (undirected) exploration by reducing the selectivity of the basal ganglia’s action selection mechanism. In contrast, exploitative behaviour was observed when reward outcomes were predictable and tonic dopamine levels correspondingly lower, enhancing the selectivity of the basal ganglia’s output. Performance was strongly determined by the range of tonic dopamine fluctuation, with best performance at intermediate dopamine levels similar to those seen experimentally (Fiorillo, Tobler, and Schultz 2003). Outside of this intermediate range, when changes in tonic dopamine where either too small or too large, decision making performance degraded, suggesting an ideal range of dopamine supports optimal explore-exploit decision making.

When combined with dopamine-dependant changes in cortico-striatal synaptic plasticity and a recent computational update to the intrinsic circuitry of the basal ganglia (Suryanarayana et al. 2019), the model could perform the multi-armed “restless” bandit task at levels comparable to that of a Kalman filter. These simulations imply that fluctuations in the level of tonic dopamine interact with the basal ganglia’s action selection circuitry to implement a biological circuit for resolving the exploration-exploitation dilemma.

## Methods

### Restless bandit task and Kalman filter

We based our 4-armed restless bandit task simulations on those described by (Daw *et al*., 2006). The task consisted of 300 trials, with the payoff for the *i*th slot machine of the bandit on trial *t* being between 1 and 100 points drawn from a Gaussian distribution (standard deviation *σ_o_* = 4) with a mean *u*_*i,t*_. At each time point the means diffused with a decaying Guassian random walk, with *u*_*i,t*+1_ = ϑ*u*_*i,t*_ +(1-*ϑ*)*θ*+*v* for each of the four choices. The decay parameter *θ* was 0.9836, the decay centre *θ*= 50 and the diffusion noise was a zero mean Gaussian with standard deviation (*σ*_*d*_)2.8.

A Kalman filter (Kalman, 1960) is the optimised mean tracking rule for this task. On trial *t* the posterior mean for the chosen option,*i* with payoff *r*_*t*_ is, 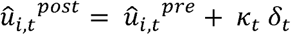 where 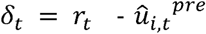, and 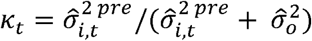 The posterior variance for the chosen option is 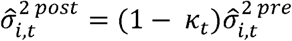 and the prior mean and variance becomes 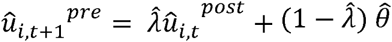 and 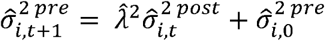 Kalman filters choices were determined by the softmax rule 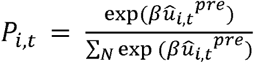. We used the parameters values from Daw et. al., (2006) as the initial starting points to optimise the Kalman filters performance of the random walk using the Matlab function *fminunc (*Mathworks, Natick). The final estimated model parameters were 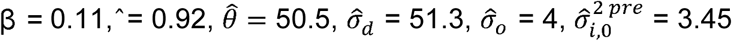 and 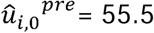.

### Basal ganglia model

The basal ganglia are a group of interconnected subcortical nuclei which receive input from most of the cerebral cortex with an output that influences the excitability of the thalamus and brainstem. Previous computational studies have demonstrated that the circuitry which comprises this structure has evolved to perform action selection. Action selection can be conceptualised as the process where by an action, *A*, (or decision) is selected from a series of *N* competing alternative choices. Within the basal ganglia, each competing action is represented by a channel, *i*, within a series of parallel re-entrant loops which run from the initial input nuclei (striatum, subthalamic nucleus) to the principal output nucleus (internal segment of the globus pallidus; GPi).

As a starting point for our simulations we used the recently described extended architecture of (Suryanarayana *et al*., 2019). This action selection circuit, which we refer to as the Gurney-Prescott-Redgrave extended model (GPRe), includes a detailed model of the basal ganglia, including the intra and inter-nuclear connectivity of the GPe (See Appendix Figure 1A). This includes details of the two subgroups of GPe cells (so-called “Inner” and “outer” populations, (Sadek *et al*., 2007) and the more recently described Arkypallidal and Prototypical GPe cells which project to the striatum and basal ganglia output nuclei respectively (Mallet *et al*., 2016).

**Figure 1:**
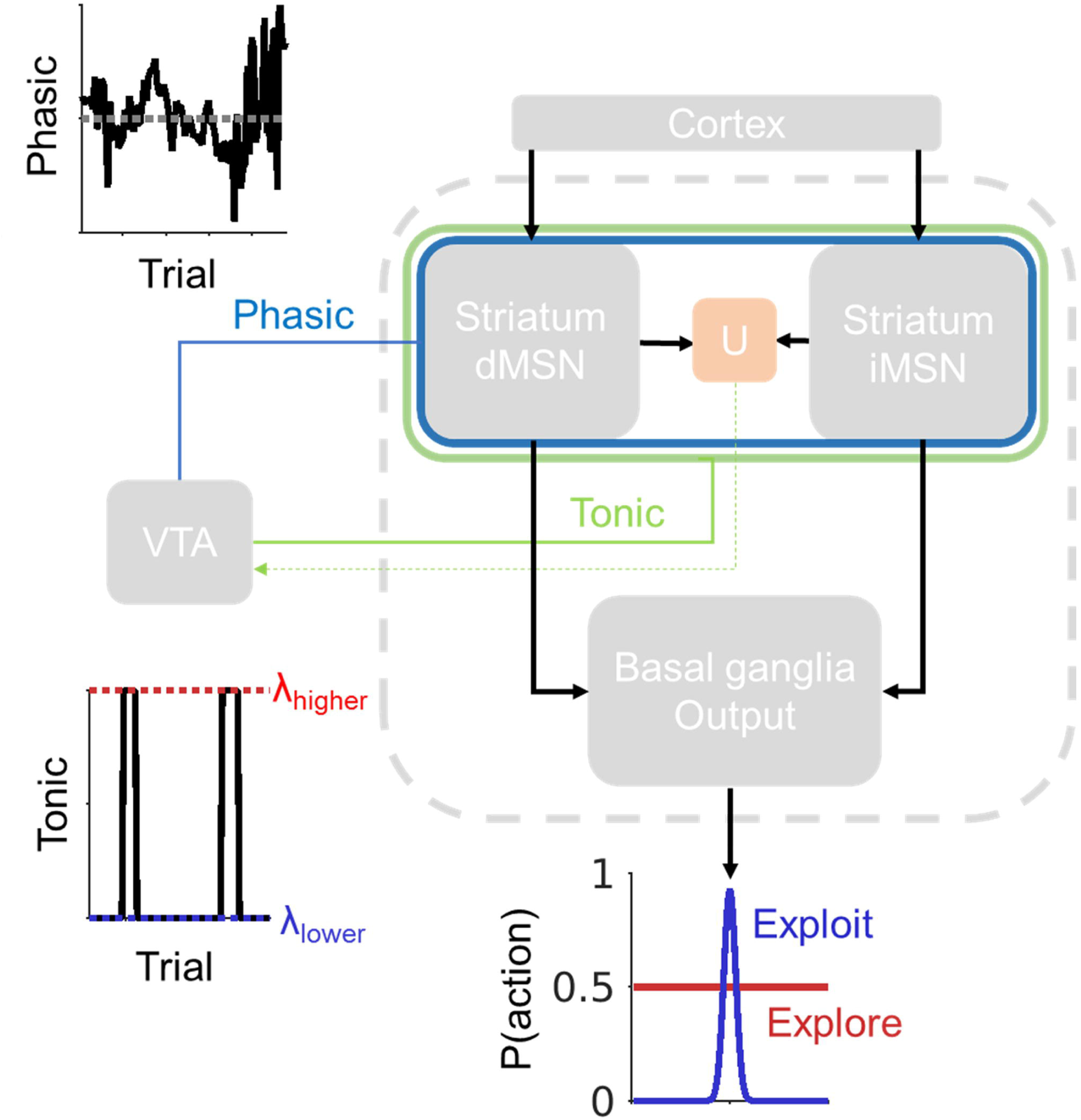
Phasic and tonic dopamine signalling influences on action selection - Model schema. Our model assumes that phasic (RPE) mediated dopamine signals are communicated to the striatal direct (D1R dominant) and indirect pathways (D2R) where they modify synaptic strength at the cortico-striatal synapse. A separate and independently regulated source of dopamine is the trial-to-trial fluctuation in the baseline tonic firing rate, which for simplicity, we assumes takes on one of two values *λ*_*higher*_ high or *λ*_*lower*_ low states, represented by the red and blue dotted lines respectively. The level of tonic dopamine is updated on each trial by a feedback from a striatal estimate of reward uncertainty (U) to the VTA (green dotted line). Accordingly, under fluctuating conditions of tonic dopamine, the corresponding changes in excitability of the select and control pathways, influence the downstream action selection function and either narrow (lower levels tonic dopamine) or expand (higher tonic dopamine) the choices available. Note that for simplicity, the influence of the tonic dopamine on phasic RPE mediated changes in cortico-striatal plasticity are not included in this illustration.

Critical to our hypothesis is the idea that the selectivity of the basal ganglia can dynamically change; switching between states of high selectivity, when there is confidence in the correct action; to low selectivity states broadening the available actions to select from (Figure 1). This is equivalent to “hard” and “soft” selection derived from classical “grid” selectivity tests (Humphries *et al*., 2006). In the simulations below we test whether these transitions in the action selection precision of the basal ganglia circuit may be a neural substrate for explore-exploit decisions.

For each trial *t* we simulate the activity of the network across a 5 second epoch with a time step of 0.1ms. For the general case *n*, the activity of the striatal neural population a in channel *I* is;

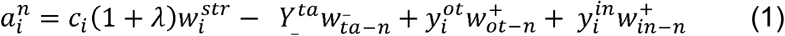

where 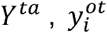, and 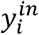 represent the activity of the GPe-Arkypallidal to striatum, the Gpe “outer” and GPe “inner” neurons respectively and 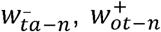 and 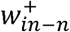 their corresponding synaptic weights. The variable *λ* here represents the level of tonic dopamine and can take one of two values (discussed below). Note that for *λ* always takes on the same value for the direct (D1R) and indirect pathways (D2R) but is always negative for the latter. For all simulations, the values of all variables as previously described (Suryanarayana *et al*., 2019) were used (see Appendix 1).

For both striatal direct 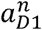 (D1R expressing) and indirect 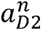 (D2R expressing) pathways, *c*_*i*_ is the cortical input to the four channels, which represents each possible choice in the restless bandit task. The salience of the cortical stimulus to each channel at each time step was fixed at a value of 0.5 modulated by uniformly distributed noise with a mean of zero and a range of ± 0.1. This was presented to both striatal populations and all channels simultaneously in time interval 1, defined as 1 ≤ t ≤ 4 seconds of the simulated trial. The selected action(s) selected were then determined by suppressing the activity of the GPi:

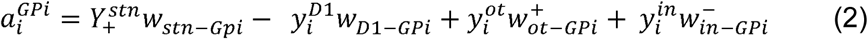

so that when its output,

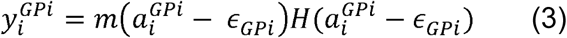

was less than the selection threshold *θ*_*d*_ (set at 0), in the time interval 2, where 2 ≤ t ≤ 3, the action in channel *i* is selected. Here *ϵ*_*GPi*_, is the output threshold of the Gpi 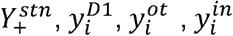,and 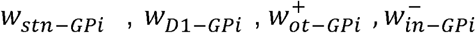 refer to output activities of the STN, direct pathway (D1R) striatum, GPe “outer” and “inner” populations and their respective synaptic weights. For the output function (3), the variable *m* is the slope of the neuronal activation function (equal to 1 for all simulations and nuclei) and *H* is a piecewise linear squashing function (Gurney *et al*., 2001). Specific details of the activation and output functions for the three GPe populations and the STN are given in Appendix 1.

Our model assumed that the background firing rate of VTA (Ventral Tegmental Area) neurons and correspondingly the extracellular striatal dopamine levels fluctuated depending upon the level of reward uncertainty. We adopted the simplest case where tonic dopamine levels fluctuate between two extremes of a range defined by *λ*_*lower*_, and upper,. *λ*_*upper*_ bounds. This approach ignores the likelihood that physiologically, tonic dopamine more likely exists across a discretely graded range, rather than a binary state, but was preferred for computational tractability at the expense of model performance. We used experimental data to approximate our initial “best guess” for the range of tonic dopamine. Recordings of primate midbrain dopaminergic neurons show a firing rate increase of ~40% when reward delivery is random (Fiorillo, Tobler, and Schultz 2003). Dopamine efflux into the NAc as measured by microdialysis in rats making forced high risk choice increase in to very similar levels (St Onge et al. 2012). Pearson et. al., (Pearson *et al*., 2009) also identified neurons in posterior cingulate cortex which increase their background firing rate from ~4 to 6Hz during explorative decision making. On the basis of these data we assumed that any increase in tonic firing rate and corresponding striatal dopamine was likely to be relatively modest and chose an initial value for *λ*_*upper*_ of a 40% greater than the lower bounds. On any trial, tonic dopamine took on one of these two values. When reward uncertaintly, *U*(*t*), was high the tonic dopamine level, *λ*, adopted the value defined by. *λ*_*upper*_ otherwise it assumed the value of *λ*_*lower*_. We further explain how *U*(*t*) is defined below.

To estimate the *λ*_*lower*_ value for the model we simulated presentation of the cortical stimuli to each of the four channels in the model across a range values of *λ* and estimated the number of channels selected by the model. We reasoned that the *λ*_*lower*_ value was best defined as the point at which the model transitioned from selecting no action, as this value when influenced by changes in synaptic weight with learning, would have the best likelihood of “hard” action selection with highest selectivity.

As the task relies on a single choice being made from the four options, when the basal ganglia co-selected more than one action, the final action chosen was determined pseudo-randomly from the available options presented. We assumed the basal ganglia model was in an exploratory “mode” at the start of the task so the initial (t = 1) value for *λ*= *λ*_*upper*_. Where values of *λ*_*higher*_ were equal to *λ*_*lower*_ no action reached the threshold for selection in either model so where not included in the analysis. The effect of the slowly fluctuating tonic dopamine level was then to *guide the precision of the action selection process* in presenting either a narrow or broad range of possible actions for any one decision.

We now describe how this can be related to the level of uncertainty *U(t)* and change in cortico-striatal synaptic weights 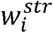 Starting with a standard delta learning rule as a model of phasic dopamine;

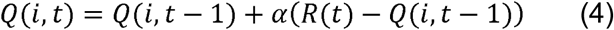

where αis the learning rate and constant for all simulations at 0.2; and R(t) is the outcome for the chosen action determined by the Gaussian random walk. The phasic dopaminergic reward prediction error (RPE) is assumed to be RPE =*(R(t) −Q(i,t −*1) and the striatal synaptic weights were updated according to:

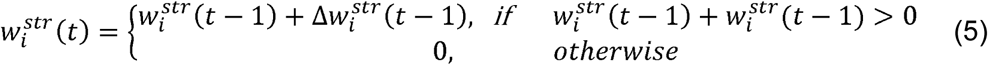

The change in synaptic weight was assumed to obey Hebbian dynamics and be the product of the pre-synaptic and post synaptic striatal activity modulated by dopamine;

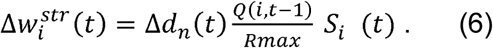

where *S*_*i*_ represents average striatal activity in time interval 2, and *Rmax*= 100. Note that we assume action-values represented by Q are computed directly in the striatum from the converging cortical inputs pre-synaptically (Samejima *et al*., 2005);

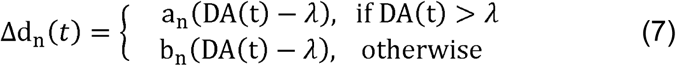

Where *a*_*n*_ and *b*_*n*_ are coefficients determining the dependence of synaptic plasticity on the current trial’s level of dopamine DA(t). Fluctuations in tonic dopamine,, also influence cortico-striatal synaptic plasticity as *λ* takes on the same values in equation (7) of *λ*_*upper*_ and *λ*_*lower*_ depending upon the level of uncertainty. As demonstrated previously (Gilbertson *et al*., 2019), for learning to occur in the model the “a” parameters (*a*_*D*1_, *a*_*D*2_) have to take positive values for the direct pathway MSNs (dMSN) with negative values for the indirect pathway MSNs (iMSN). This is consistent with the opposing effects of the positive prediction error signal on D1 and D2 receptors via LTP and LTD. Conversely, the *b* parameters (*b*_*D*1_, *b*_*D*2_,) which govern the magnitude of LTD and LTP at the direct and control indirect pathway synapses, must also have negative or positive values respectively. Equation 7 links the RPE from Equation 4 by;

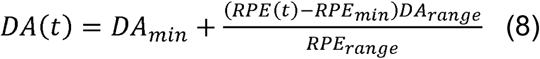

where RPE(t) < 0, *DA*_*min*_= 0, *DA*_*range*_= *RPE*_*min*_ = -1, *RPE*_*range*_ = 1; otherwise *DA*_*min*_ =*DA*_*range*_ = 1 - *RPE*_*min*_= 0, *RPE*_*min*_ = 1.

We also top-limited the weight changes of both sets of cortico-striatal synapses (direct and indirect) to prevent saturation of the striatal activity so that;

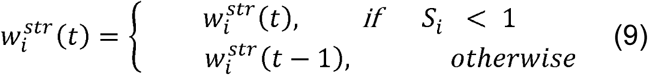

Equally, when the selection threshold of the activity in the GPi is not met and no action is presented, the synaptic weights, Q-values and RPE signals are not updated and remain the same as the previous trial. For all simulations we assume that the initial Q values for all actions for trial 1 are set to 50 and the striatal synaptic weights for both the direct and indirect pathways were initialised at 1.

The basic premise of our model is that when the identity of the highest value action is unknown, a *feedback loop* between a neural estimate of this uncertainty triggers an increase in background dopamine levels. This, in turn, softens the action selection mechanism and promotes random exploration of alternative actions. One potential source of this uncertainty signal is encoding by the striatal weights of the direct and indirect pathways (Mikhael & Bogacz, 2016). We found the most robust approach to identify “transition” points in the random walk (i.e. points where a previously high value choices value decays and is replaced by one of the other three options), was to calculate our uncertainty index, *U(t)*, using the difference in the weight values between the direct and indirect pathways. The utility of this signal can be understood be considering the effect of prediction error signalling on the direct and indirect pathway weights during a transition in value between actions. As the value of a previously high value action decays the negative phasic prediction error signal associated with lower than expected outcome would be expected to decrease the direct pathway weights whilst increasing the indirect pathways weights. Until a novel, alternative action is “explored,” and its outcomes deemed suitable to “exploit”; the relative difference in weights for that channel 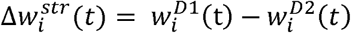 tend towards zero and values less than zero. This means that when all the 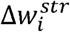 values for all four options are close to or less than zero, uncertainty about the highest value choice is high.

Again, here we execute the most tractable solution computationally, whilst acknowledging its biological simplicity, where uncertainty exists in a binary state and is either a high *U(t)*= 1 or low *U(t)*=0 value. We then map the level of tonic dopamine directly unto the level of uncertainty so that when (*t*)= 1, *λ* (*t*) = *λ*_*higher*_, *otherwise λ(t)*= *λ*_*lower*_. Empirically, we found that model performance was dependent on the addition of a threshold,*U*_*thres*_, which determined when a difference in the synaptic weights was worthy of a transition in the uncertainty level. Accordingly, we define the uncertainty *U(t)* in relation to this threshold, *U*_*thres*_ by:

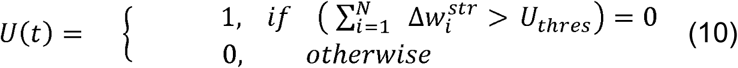

If *U*_*thres*_ is set too close to zero, a small increase in the 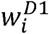 synaptic weight for one action meets the criteria for values of *U(t)*= 0 and correspondingly lower values of tonic dopamine set by *λ*_*lower*_. If this increase in the direct pathway weights is not sufficiently large to overcome the akinetic effects of tonic dopamine at its lower bounds, *λ*_*lower*_ no action is disinhibited by the basal ganglia’s output nuclei, and the model is unable to perform the task. We used a constant value of *U*_*thres*_ = 0.1 for all simulations. This allowed the 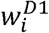 synaptic weights to build sufficiently enough for the action selection circuit to disinhibit a single action for selection, when the tonic levels of dopamine were at the lower end of their range.

#### GPR model

The original GPR model (Gurney *et al*., 2001) consisted of the same basic control and select pathways as the extended (GPRe) version but without the additional intrinsic and extrinsic connectivity of the GPe. Our rational behind simulating the effects of slowly fluctuating tonic dopamine on this model was to establish that the additional biological detail in the GPRe action selection mechanism was necessary for optimal performance of the task. The equations pertaining to the activation and output functions for the GPR model are included in Appendix 1. The schema for the GPR model is also included in Appendix Figure 1B.

### GPR model

The original GPR model (Gurney *et al*., 2001) consisted of the same basic control and select pathways as the extended (GPRe) version but without the additional intrinsic and extrinsic connectivity of the GPe. Our rational behind simulating the effects of slowly fluctuating tonic dopamine on this model was to establish that the additional biological detail in the GPRe action selection mechanism was necessary for optimal performance of the task. The equations pertaining to the activation and output functions for the GPR model are included in Appendix 1. The schema for the GPR model is also included in Appendix Figure 1B.

## Results

### Defining the dynamic range of tonic dopamine and cortico-striatal plasticity in the model

Before assessing the basal ganglia model’s performance on the task, we performed a series of simulations to define the range of values that the tonic level of dopamine would take. Defining the lower bounds of the tonic dopamine range, *λ*_*lower*_ was of particular significance. This value, in combination with the influence of learning induced changes in synaptic plasticity, would determine how “hard” the selection mechanism of the basal ganglia was during periods of low uncertainty. It is during these periods that we hypothesise that exploitative choices are sustained by highly selective and precise action selection. Previous simulations have emphasised a narrow range of tonic dopamine that can produce “hard,” action selection, with values above or below this optimal level leading to significant softening (multichannel activation) or non-selection of actions (Suryanarayana et al. 2019). In classical “grid” based tests (Humphries *et al*., 2006), which examine the influence of dopamine on the selectivity of the action selection circuit, cortical stimuli are presented sequentially, to a pair of channels, across a range of stimulus saliencies (intensities). In our case, the cortical stimulus had a fixed salience and was presented to all channels in the model simultaneously, to emulate the presentation of the four choices available in the task. We simulated presenting this stimulus at a series of dopamine levels within the range ∈ [0.15, 0.6]. Each simulation was run 100 times and the number of channels *N*_*c*_, selected by the model estimated (Figure 2E).

**Figure 2:**
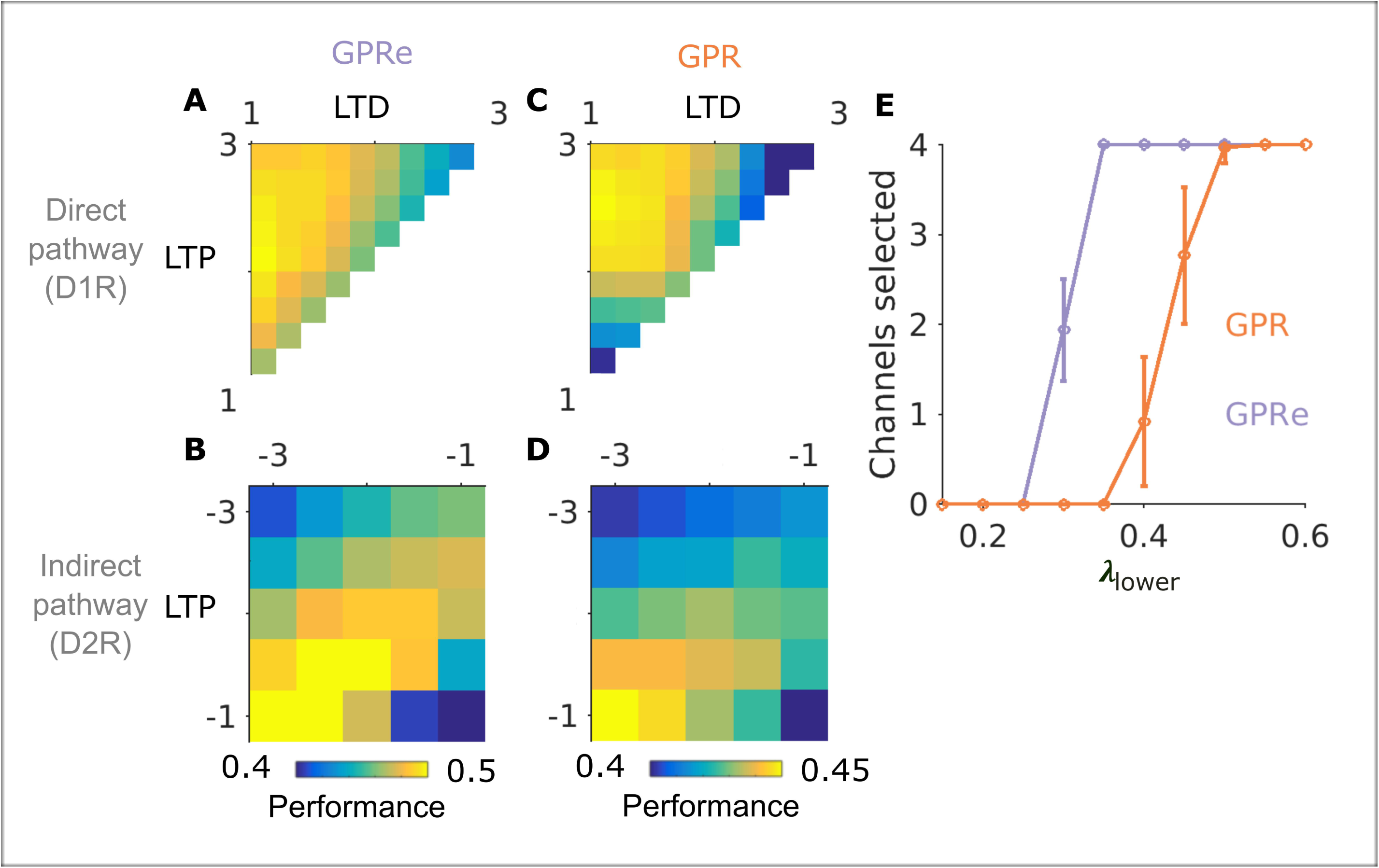
Defining the striatal plasticity parameters and optimal range of tonic dopamine. Colour maps representing the GPRe and GPR action selection circuits performance 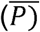 in the restless bandit task for different values of D1 and D2R cortico-striatal plasticity parameters. Task performance was best when D1-LTP was significant greater than D1-LTD as no action-value acquisition occurred when LTD dominant (**A** – GPRe, **C** – GPR). In contrast, optimal task performance required LTD to be greater than LTP cortico-striatal plasticity at D2R in the control pathway (B-GPRe, D, GPR). Models were D2R-LTP was stronger than LTD led to a build-up of striato-pallidal activity which led to akinesis and no action being dis-inhibited by the basal ganglia’s output nuclei. (E) The effect of varying the level of lowed bounds for tonic dopamine (*λ*_*lower*_) on the average number of actions disinhibited (selected) in the basal ganglia’s output nucleus (GPi) in the two action selection circuits (GPR-red) GPRe (blue). Error bars represent Standard deviation.

Consistent with the effects of dopamine on the model’s excitability, low values of *λ*, corresponding to low tonic dopamine lead to *akinesia*, where no action is selected. In contrast, as *λ* increases, the balance of excitability within the direct and indirect pathways leads to the selection of one or more actions. We reasoned that the transition point, that being the highest value of that produces akinesia, would represent the lower bound for tonic dopamine (*λ*_*low*_). Although this value did not generate any actions, we expected with learning and the associated changes in synaptic weights, that this lower bound would be associated with the highest levels of selectivity, necessary for single action selection.

Following this procedure, the *λ*_*lower*_ value was set at 0.25 for the GPRe and 0.35 for the GPR model. Setting the upper range (*λ*_*higher*_) of dopamine at +40% of this was based on physiological studies (Fiorillo *et al*., 2003; St Onge *et al*., 2012), *λ*_*higher*_ for the GPRe and GPR models was set at 0.35 and 0.5.

Next we mapped the parameter space of the four variables in the model which govern cortico-striatal plasticity. The motivation behind this was to maximise the models performance in the task by minimising the playoff between the synaptic weight changes on the excitability of the action selection circuit. This is because the synaptic weights contribute to the overall excitability of the direct and indirect pathways and the balance of their excitability determine whether or not an action is selected. For example, if the excitability changes induced synaptically by learning in the direct to indirect pathway are not suitably balanced, the models performance will be artificially penalised trials were no action is selected. For the purpose of this mapping procedure, we placed several constraints on the bounds of the parameters. First, we assumed positive values for the parameters pertaining to cortico – striatal plasticity at D1R expressing striatal neurons (*a*_*s*_, *b*_*s*_) and negative values for those relating to plasticity at D2R expressing neurons (*a*_*c*_, *b*_*c*_). Not only does this significantly eliminate redundancy in the parameter space (Gilbertson et. al., 2019) but it makes physiological sense, as these govern the gradient of the dopamine-weight change curve for both subclasses of receptors (Figure 2). In addition, for the D1R parameters we did not include any combination of parameters where (*a*_*s*_ > *b*_*s*_) as the makes D1-LTD>LTP and no learning is possible. For further computational tractability we limited our parameter search to an interval of [1,3] [1,3],[-3, -1],[-3, -1] with steps of 0.5 for *a*_*s*_, *b*_*s*_, *a*_*c*_, *b*_*c*_ respectively, including every possible combination within this range.

As a performance index we defined the probability of making the highest value choice through the task 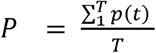 where, T is the number of trials, (n= 300) and 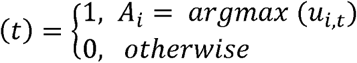,*u*_*i,t*_ is the payout for all bandits at trial t. A *P* value of 1 reflects perfect choices were the highest value choice is made on every trial. We estimated values for each plasticity combination after 100 runs of the task to produce an average performance index 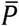. For all iterations, the same Gaussian random walk was presented to the model on each run. For both the GPRe and GPR selection mechanisms, this converged on the same combination of direct (D1R) and indirect (D2R) plasticity parameters where; *a*_*s*_ = 1, *b*_*s*_ 1.5, *a*_*c*_ = -1.5, *b*_*c*_ -2.5 with a value of 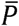 of 0.54±0.04, 0.47±0.03 respectively (Figure 2A & B for GPRe; C&D correspond to the GPR mapping). A similar analysis of performance applied to the Kalman filter solution to the same random walk produced a *P* value of 0.58.

### Model performance in the restless-bandit task

Figure 3 is an illustrative example of both the Kalman filters (B-D) and the GPRe models choices (E-F) from a representative single run (GPRe model *P* = 0.66). During transition points in the Gaussian random walk, where the value of a previously high value action decays and is replaced by an another action, the Kalman filter “exploration” of alternatives correlates with an transient increase in the Kalman gain *κ*. The basal ganglia’s choices are driven by the level of tonic dopamine (Figure 3G) and its influence on the selectivity of the basal ganglia’s action selection mechanism (Figure 3H). Here, selectivity *S* is defined as 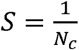, where *N*_*c*_ is equal to the number of channels selected (disinhibited) by the GPi (values of <1 reflect more than 1 channel is selected). Consistent with their opposing effects on the basal ganglia’s output nucleus’s excitability, the synaptic weights increase in the direct (D1R) and decrease in the indirect (D2R) pathways as the higher value choice is identified (Figure 3I). In contrast, the cortico-striatal weights decrease in the direct pathway for as the value of the action decays in line with the random walk, and the indirect pathway weights increase to values >1, if a choice leads to a below average (<50) payout (see choice represented by blue line Figure 3I, direct pathway weight from trial 180 onwards). In Figure 3J, the trial-to-trial difference in synaptic weights between the direct and indirect pathways is used as an estimate of the uncertainty *U*(*t*) which we proposes feeds back to govern the background level of tonic dopamine (Figure 1 model schema).

**Figure 3:**
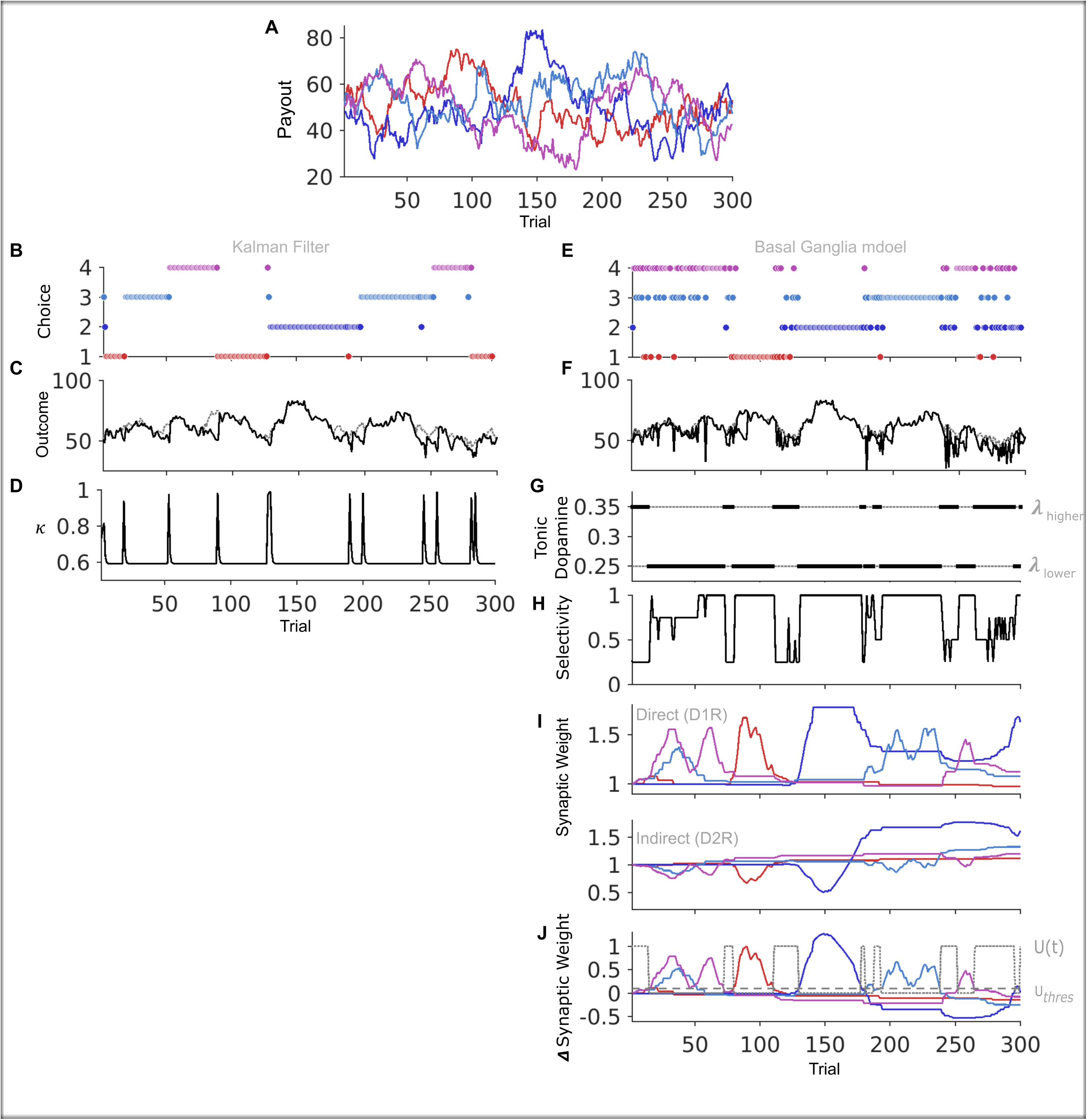
Comparison of basal ganglia model and Kalman filter performing the restless bandit task on a single run. **(A)** Payout of the Gaussian random walk for each of the four “bandits”. (B) The kalman filters trial-to-trial choices, (C) actual payout (black line), ideal payout(grey line) led to the probability of choosing the highest value bandit *P* = 0.58. We also plot the kalman gain (*K*) in (D). An example of the the full basal ganglia model (GPRe selection mechanism) performing a single “run” of the task is illustrated for comparison. With the basal ganglia models choices (E), payout (F) and the corresponding trial-to-trial modulation in the tonic dopamine level (G). Here this is constrained by the upper and lower bounds of the background dopamine ranged defined by *λ*._*higher*_ and *λ*._*lower*_ respectively. The influence on the models action selection “selectivity”, defined as the inverse of the number of actions disinhibited by the basal ganglia’s target nuclei. Values of 1 represent single “hard” action selection trials, whereas during trials were this is 0.25, “soft” action selection leads to four actions being dis-inhibited allowing random exploration of available options. The cortico-striatal synaptic weight for each action in both the direct and indirect pathways are plotted in (I) with the derivation of the level of Uncertainty, *U*(*t*) represented by the grey dashed line in (J). This is calculated from the difference in the synaptic weights, and thresholded by the value of *Uthres*. The influence of the level of uncertainty on the tonic dopamine can be seen by comparing the time course of changes in *U*(*t*) with the tonic dopamine level in (G). On this run the basal ganglia’s probability of choosing the highest value action was *P* = 0.66.

In Figure 4, we illustrate the average performance across multiple simulations (n=100), including the trial – trial change in choice probability (Figure 4B), tonic dopamine, (Figure 4D) and average selectivity (Figure 4E) in the GPRe model. The effect of fluctuating levels of tonic dopamine then become apparent on the action selection mechanism of the basal ganglia. Under conditions of high dopamine, for example, at the beginning of the task, selectivity is low, all channels are disinhibited by the GPi, and all four choices are available to “explore” their value. In contrast, once the correct choice is established, the levels of dopamine drop to the lower end of its range (*λ*_*lower*_), and a single action is presented, allowing this newly discovered, high value choice to be “exploited”. Because synaptic level changes evolve slowly between trials, these become an unreliable means to make decisions when there are rapid changes in the reward environment (Drummond & Niv, 2020). This inflexibility is overcome by more rapid fluctuations to higher tonic dopamine states which accompany the transition points in the random walk, where one choices value diminishes and another rises with the random diffusion process. Here, the role of the synaptic weights is in estimating the uncertainty of the reward schedule (Equation 10) and feeding this information back to determine the background levels of dopamine. The behaviour of the model becomes a constant *feedback loop* between three interacting functions: 1) phasic dopamine’s influence on the striatal weights through the RPE: 2) intra-striatal uncertainty estimate and finally: 3) effect of this on the level of tonic dopamine and its influence on the selectivity of the basal ganglia action selection circuit.

**Figure 4:**
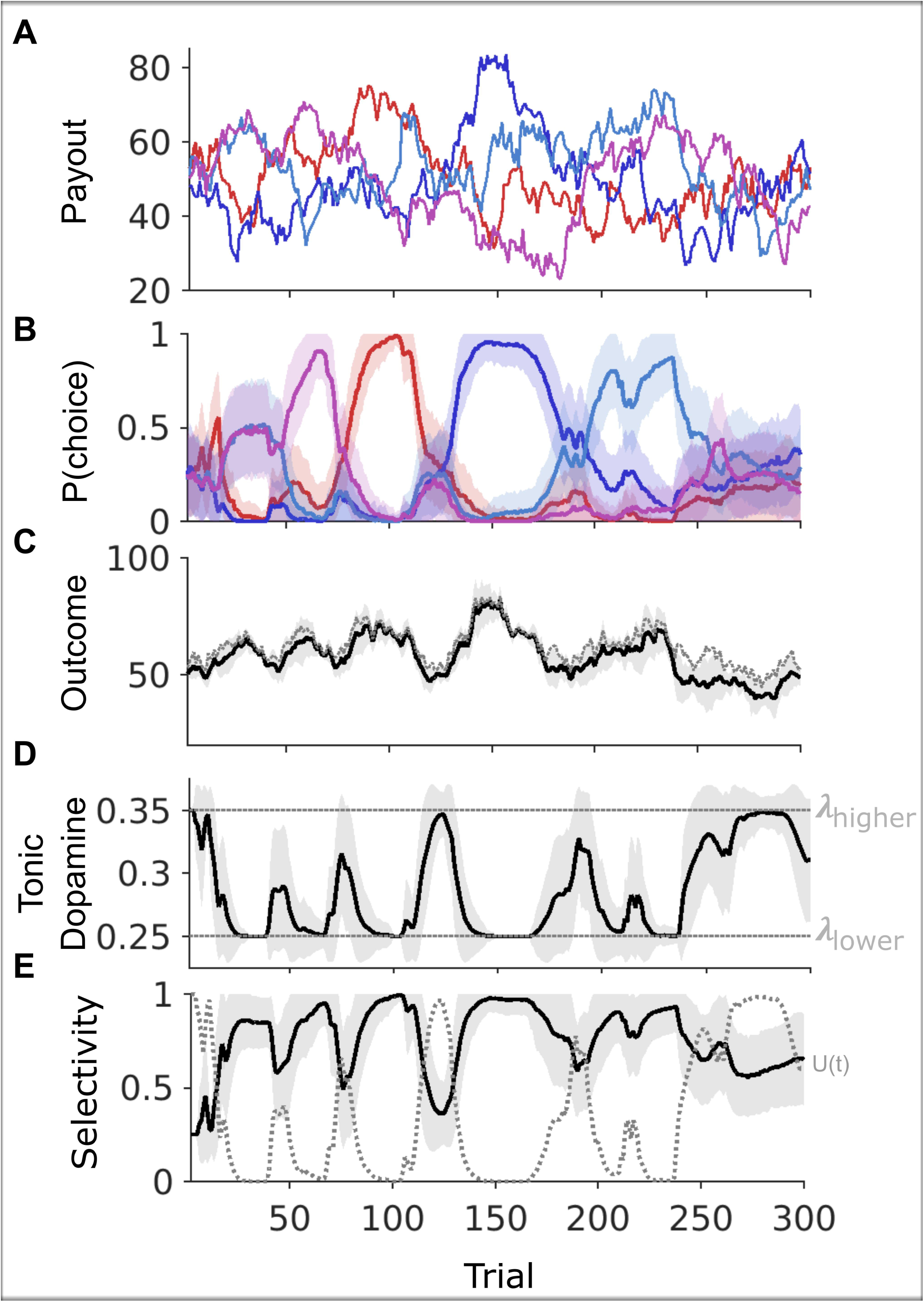
Average performance of the basal ganglia model performing the restless bandit task. **(A)** Payout of the Gaussian random walk. (B) Choice probability (B), payout (C), trial-to-trial change s in tonic dopamine (D) and the selectivity of the action selection circuit (E). All values derived from averaging (n=100) simulations of model performing the same random walk. Shaded regions represent standard deviation. Selectivity was calculated by taking the reciprocal of the number of actions (channels) disinhibited by the model GPi. The dotted grey line in (E) represents the uncertainty estimate which determines the tonic dopamine levels.

### Influence of action selection mechanism

Next, we explored the influence of the action selection mechanism on the models performance. Recall that by including additional intra- and inter-nuclear GPe connections, the action selection function is enhanced in the GPRe model, Suryanarayana et. al., (2019), compared to the GPR model (see also (Bogacz *et al*., 2016) for evidence in support of these additional connections enhancing action selection function). We predicted that if the action selection mechanism is critical to decisions in the restless bandit task, this should lead to heightened performance by the GPRe model. We ran a series of simulations comparing both model’s performance across a range of tonic dopamine. Here, the lower bound of the tonic dopamine was kept constant for both models, as were all other parameters. The upper bound (*λ*_*higher*_) for the tonic dopamine was varied to levels of *λ*_*lower*_ 20, 40, 60, 80 and 100% of its value (Figure 5).

**Figure 5:**
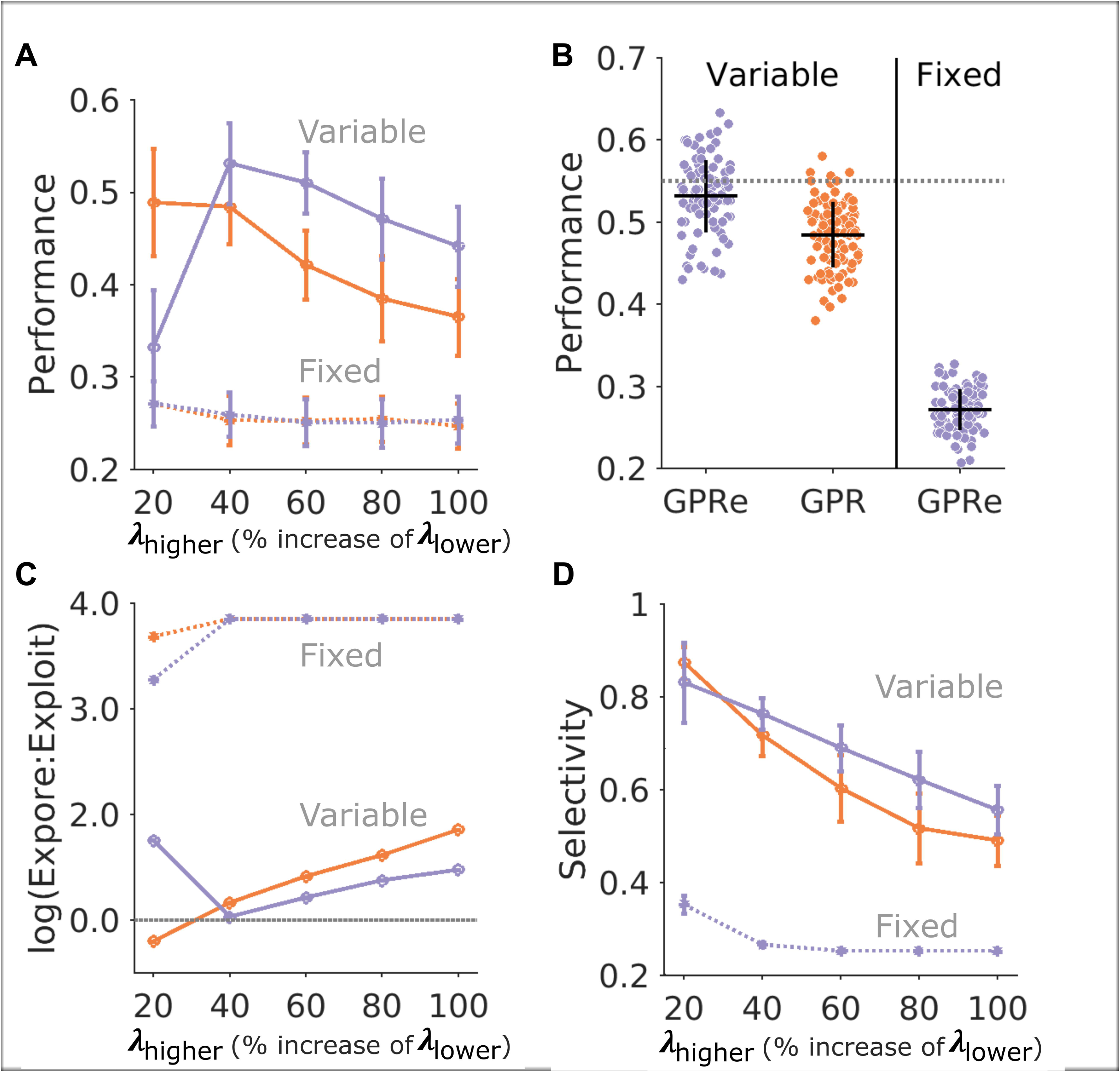
The effects of the action selection mechanism and tonic dopamine on model performance. (A) Varying the levels of the upper bound, *λ*_*higher*_, of tonic dopamine (here expressed as a percentage increase of the lower bound) leads to optimal performance of the GPRe selection mechanism at +40% (Red line GPR, blue line GPRe). Constraining the tonic dopamine levels to a “fixed” value, rather than allowing this to fluctuate in a variable, trial-to-trial, basis leads to a significant deterioration in performance (dotted lines represent simulation results with “fixed” tonic dopamine). (B) Scatter plots for each simulation runs performance values. Cross hairs represent mean and standard deviation. Grey dotted line is the Kalman filters performance level plotted for comparison. (C) The E ratio (logarithm of the ratio between number of trials with a single action selected to the number of trials with more than 1 action selected) and Selectivity (reciprocal of number of actions selected) values for each model and tonic dopamine levels studied. All points represent average ± standard deviation (n=100 simulations).

Using the average performance metric 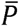 as the variable of interest, and the model type and (GPRe, GPR) and *λ*_*higher*_ as the independant variables, a two-way ANOVA demonstrated a significant effect of both model F(1) = 96.3, p<0.001 and *λ*_*higher*_ F(4) = 144.1, P<0.001. A direct comparison between the GPR and GPRe performance demonstrated that the GPRe performance was consistently better than the GPR (*λ*_*higher*_ = *λ*_*lower*_ + 40%; 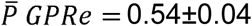, *P* GPR = 0.47±0.03, paired T-test, t (198) = 5.9, p< 0.001). This enhanced performance was also associated with greater average selectivity, 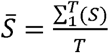 across all ranges of *λ*_*higher*_ in the GPRe model, 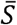 GPRe = 0.69±0.03, GPR = 0.64±0.04, with a significant effect of the action selection model (Two-way ANOVA F(1) = 197.3 p<0.001). 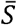 values closer to 1 indexed how consistently the action selection circuit selected a single channel from the four options available.

As a metric of how likely the model selected multiple choices during periods of high uncertainty, we defined the exploratory ratio as, 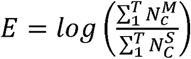, where 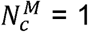 when *N*_*c*_ (*t*) > 1and 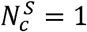 otherwise. For ratio values close to 0 the model chooses equal numbers of trials where either single or multiple channels are selected by the action selection circuit. For values of E > 0, multi-channel selection (i.e > 1) are favoured. Figure 5C illustrates the influence of the upper limit of tonic dopamine on E. Correlating with the “inverted -U” shaped effect of *λ*_*high*_ on the GPRe models performance (Figure 5A), the E value was closest to 0, at *λ*_*high*_ = *λ*_*low*_ + 40%. The model’s peak performance at this level of tonic dopamine was associated with an equal number of trials where *a* single channel was selected and trials where multiple channels were selected, supporting a balanced strategy of both exploration and exploitation.

### Effects of fixed versus fluctuating tonic dopamine levels

To further confirm that fluctuating tonic dopamine, rather than its absolute level determined performance, we re-ran the simulations but fixed the level of *λ*_*higher*_ so that it was fixed to the

same values as *λ*_*lower*_. Despite all other components of the models remaining the same, both the GPRe and GPR variants were reduced to chance levels of performance 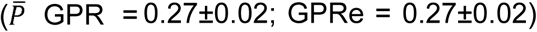 when tonic dopamine levels were equal for values across the same a range of values (Figure 5A&B). A Two-way ANOVA with independent variables including model type (GPR, GPRe), *λ* range (fixed, variable) and *λ*_*higher*_ (*λ*_*lower*_ 20, 40, 60, 80 and 100%) confirmed this significant effect of fluctuating tonic dopamine levels on performance (F(1) = 81.6, p<0.001).

Tonic dopamine exerts two effects on the model, both on the action selection mechanism and also at cortico-striatal synapses. For baseline levels of tonic dopamine, the range over which phasic increases (positive prediction error signal) influence both D1 and D2R plasticity far exceeds that for phasic reductions below this (Gurney *et al*., 2015). In principle therefore, for higher levels of tonic dopamine, the range over which the negative prediction error signal exerts its influence should be greater. To test this possibility we estimated the mean synaptic weights at each trial for both high and low levels of tonic dopamine. We also further analysed trials according to whether the reward prediction error was positive or negative in order to look for any asymmetry in the influence of tonic dopamine on RPE mediated synaptic plasticity. Performing an ANOVA with the synaptic weight taken from the GPRe model simulations as the variable of interest, we including three independent variables: Tonic dopamine, (*λ*_*higher*_, *λ*_*lower*_; dopamine receptor type, (D1R, D2R); and RPE sign, (positive or negative). This demonstrated a significant interaction between the tonic dopamine level and the dopamine receptor type (F(1) = 472.0 p<0.001, but no additional interaction or effect of the RPE sign (F(1) = 0.12, p=0.72). On average, synaptic weights at dMSN (D1R expressing) were more likely to strengthen under low levels of tonic dopamine (mean dMSN weight *λ*_*higher*_ = 1.12±0.06, mean dMSN weight *λ*_*low*_ = 1.17±0.04, T(398) = 9.77, p<0.001), whereas high tonic dopamine led to greater cortico-striatal synaptic potentiation at D2R expressing iMSNs (mean iMSN weight *λ*_*high*_ = 1.22±0.08, mean iMSN weight *λ*_*low*_ = 1.09±0.05, T(398) = 19.6, p<0.001). This analysis illustrated in Figure 6 suggested that fluctuating levels of tonic dopamine could mediate their effect on decision making by modifying striatal value estimates represented in the cortico-striatal synaptic weights.

**Figure 6:**
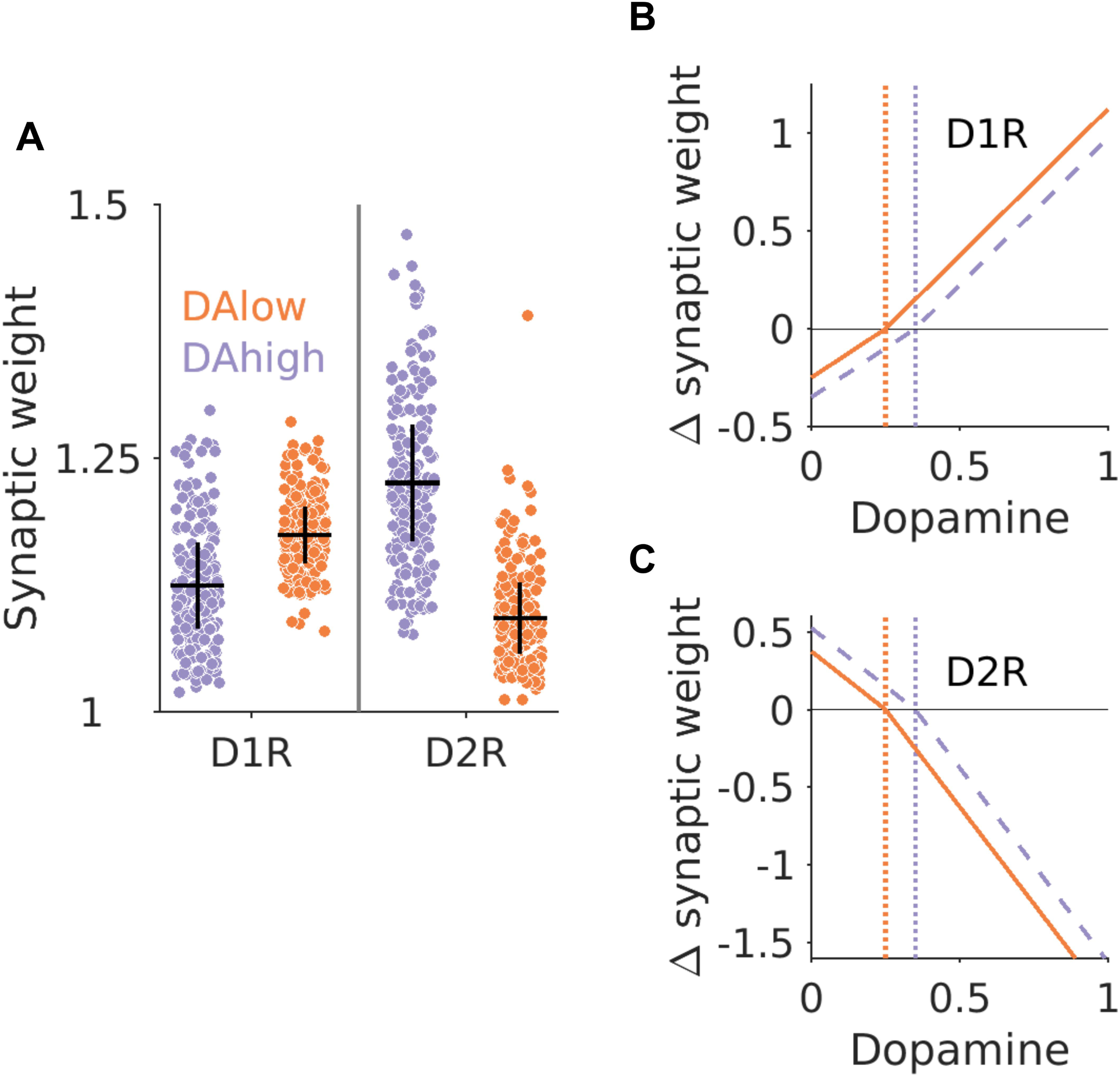
Influences of Tonic dopamine on cortico-striatal plasticity. (A) Synaptic weight changes at dMSN (D1R expressing) cortico-striatal synapses are greater when tonic dopamine levels are at the lower end (*λ*_*lower*_) of the optimal range than at the optimal upper limit (*λ*_*higher*_). This effect is predicted by the dMSN dopamine weight-change curve illustrated in (B). Here the blue vertical dotted line represents (*λ*_*higher*_), the red vertical line (*λ*_*lower*_). For dMSNs, phasic increases (positive prediction error) in dopamine above these values produce LTP and increase the synaptic weight. The effect of lowering the level of tonic dopamine to values of *λ*_*lower*_ is to amplilfy the influence of positive prediction error (phasic increases) over phasic reductions in signals. The converse is due in iMSN neurons where *λ*_*higher*_ values generate greater phasic pauses in dopamine (relative to the tonic level) which promote LTP in the iMSN cortico-striatal synapse (C).

To establish the extent to which this was important for the model’s performance of the task, we compared the original GPRe model with two further model variations (Figure 7). In the first variation (‘noplast’), we allowed the fluctuating levels of tonic dopamine to influence the action selection circuit but assumed a constant level of dopamine influenced cortico-striatal plasticity. This was implemented by allowing *λ* in equation (1) to vary according to the bounds set by *λ*_*higher*_ and *λ*_*lower*_ whilst keeping the value of that influenced synaptic plasticity in equation (7) constant between trials at the value of *λ*_*higher*_. For *λ*_*higher*_ levels set at *λ*_*lower*_ + 40%, where performance of the GPRe model was optimal 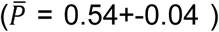, the equivalent model with no influence of tonic dopamine on plasticity, ‘noplast’-GPRe, performance was inferior, 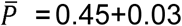 (Two tailed T-Test(198) = 3.14, p = 0.002), as it was across the same range of tonic dopamine values tested previously (One-way ANOVA F(1) = 450.25, p<0.001). We then compared these simulations to a final model variation where cortico-striatal plasticity changes were modulated by fluctuating levels of tonic dopamine but the action selection circuit was not (‘noAS’-GPRe). This was achieved by fixing the tonic level of dopamine (*λ*) to values of *λ*_*high*_ in the action selection circuit (Equation 1), whilst allowing this to fluctuate between *λ*_*high*_ and *λ*_*low*_ at the cortico-striatal synapse (Equation 7). The effect of this was to degrade the models performance to chance levels, across all values of tonic dopamine tested, including the optimal range of tonic dopamine for the full model (‘noAS’-GPRe *λ*_*high*_ = *λ*_*low*_ + 40%, 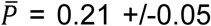 (Two tailed T-Test(198) = 16.54, p < 0.001).

**Figure 7:**
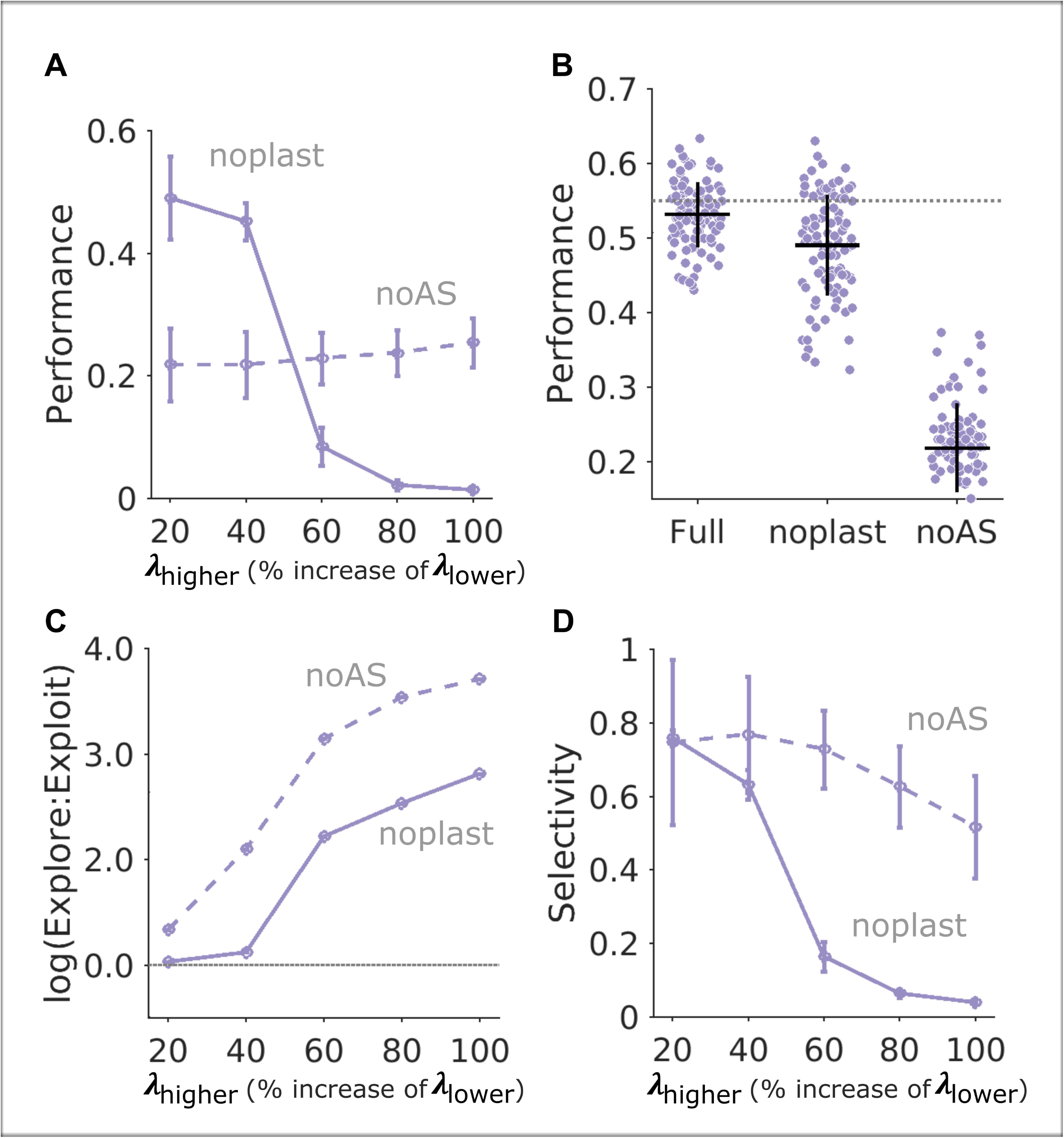
Tonic dopamine influences model performance at both the level of action selection circuit and at the cortico-striatal synapse. Here we examine the GPRe model under two further variations. The solid blue line represent the GPRe models task performance with the optimal levels of fluctuating tonic dopamine intact but allowing this to influence the action selection circuit in isolation (A). Here the “noplast” variation of the model assumes a constant level of tonic dopamine at the cortico-striatal synapse. The performance of this variation is inferior to the Kalman filter, grey dotted line in (B), significantly better then fluctuating dopamine is restored at the cortico-striatal synapse withdrawn from the action selection circuit (‘noAS’), represented by the dashed blue line. The corresponding variations in model performance are reflected in their E ratio and selectivity values. All points represent average ± standard deviation (n=100 simulations).

Finally, to examine the relationship between the action selection circuit’s selectivity and the performance of the task across the different model variations we correlated the average selectivity and E ratios with each models performance value 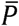 (Figure 8). Consistent with the critical role of the effect of tonic dopamine on the action selection circuit in determining the explore-exploit strategy, model performance 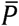 correlated significantly with both the average selectivity for the model 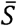 (rho = 0.77, p< 0.001) and E ratios (rho = 0.64, p<0.001).

**Figure 8:**
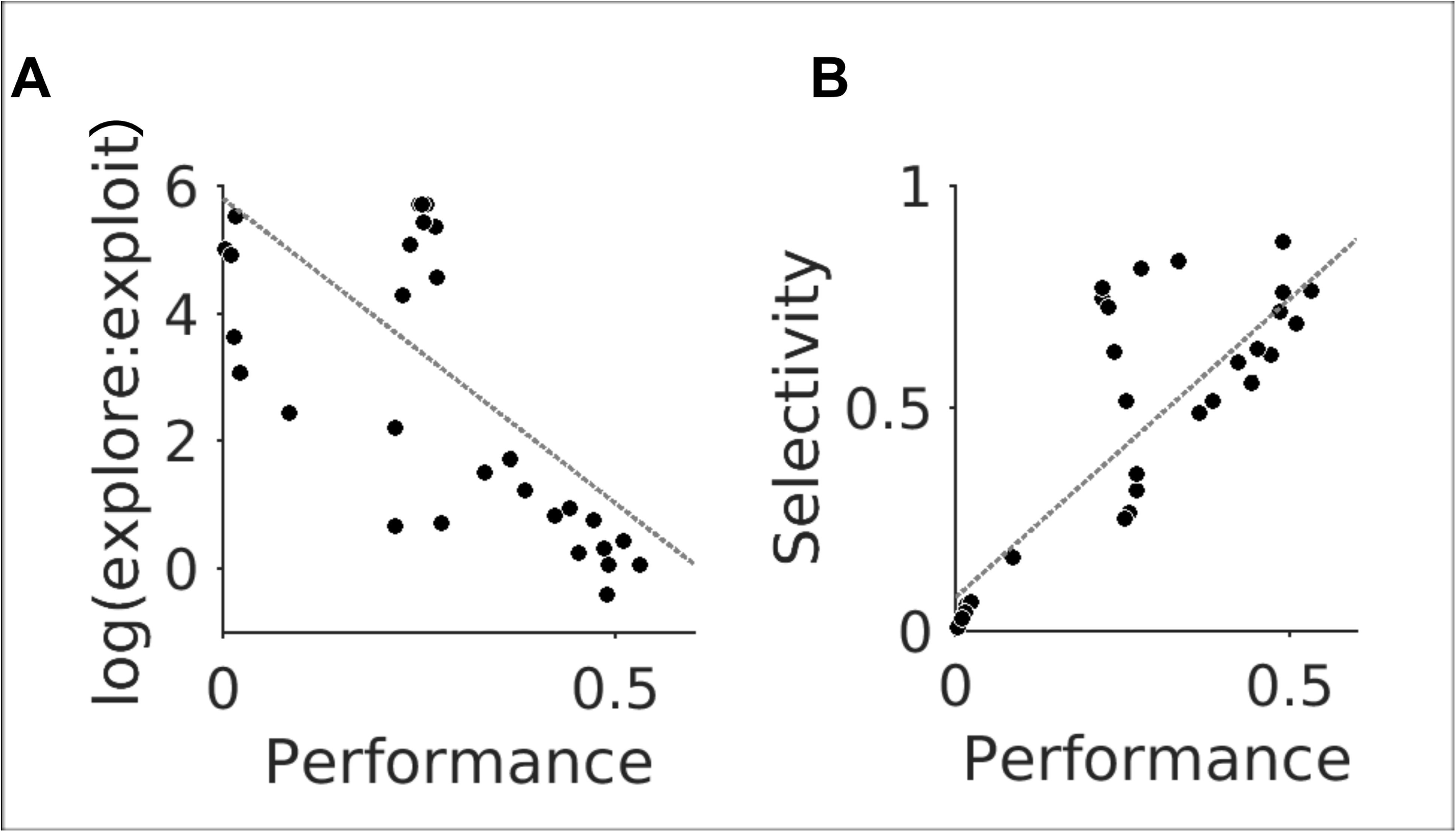
Correlations with Model performance. Across all of the model variations the selectivity (B) and E ratio indices correlated significantly with the performance in the task with the optimal performing model having both high selectivity (A) (rho = 0.77, p< 0.001) and a E ratio of close to 0 (rho = 0.64, p<0.001), consistent with a balanced decision making strategy between exploration and exploitation.

## Discussion

Making good decisions requires a trade off in the way we exploit information whilst also being amenable to explore alternative strategies. Given the significance of this problem to survival, and its relevance to diseased states decision making, understanding its neural basis has been the subject of significant past and ongoing debate. Here we propose a theoretical account of how the basal ganglia circuitry, in isolation, can provide an optimal solution to explore-exploit “dilemma”. Our model builds on previous computational studies which have argued that the extensive inter- and intra-nuclear connectivity of the basal ganglia’s circuitry, has evolved to perform the function of action selection. In the context of explore-exploit decision making, this function sub-serves a delicate balance to allow optimal behavioural flexibility. On one hand, the basal ganglia’s output can limit the set of choices when confidence is high that a choice is reliable enough to “exploit”. Equally, when the reward environment is unpredictable, its output ‘acknowledges’ this uncertainty by relinquishing control over a single action and facilitating “exploration” of alternative choices. In our model, the currency which determined these transitions in selectivity was the level of tonic dopamine. The idea that this might govern explore-exploit decisions by modulating excitability of the basal ganglia’s output is not new (Humphries *et al*., 2012). The basal ganglia has also been proposed as a circuit for explorative decision making (Sheth *et al*., 2011; Kalva *et al*., 2012; Costa *et al*., 2019).

The significance of our results is that they demonstrate that the basal ganglia is endowed with the apparatus capable of optimally tracking a non-stationary reward environment. The neural basis of how learning occurs under volatile environments is less well understood than learning in more classical stationary contexts. A pervasive view is that although the basal ganglia can excel at habitual (RPE-mediated) learning, and are likely to ‘co-operate’ via extensive re-entrant loops with the frontal cortices, in goal directed learning, but are subservient to top-down cortical influences under conditions of greater uncertainty (Daw *et al*., 2005; Cohen *et al*., 2007). Empirical evidence from functional imaging studies showing activation of prefrontal cortical regions during exploratory behaviour which have recently been shown to be dopamine dependant (Daw *et al*., 2006; Chakroun *et al*., 2020) would support this view. Neural circuits designed to approximate the function of the Kalman filter, when applied to decision making, also assume top down influence from cortical centres such as orbitofrontal cortex on the striatum (Gershman, 2017).

Here we propose that the basal ganglia has a far greater *intrinsic* repertoire of algorithmic solutions to learning than isolated RPE “model-free” learning. One possible example of this enriched repertoire, is combining classical RPE signalling with dynamic fluctuations in tonic dopamine. This is an integrative account of phasic-tonic learning signals aligns itself to the growing theoretical evidence re-evaluating the computational accounts of more sophisticated intrinsic processing within the basal ganglia (Mikhael & Bogacz, 2016; Dunovan *et al*., 2019; Bogacz, 2020). The implication of our results is that that the basal ganglia may, alongside its prefrontal cortical brethren, provide one of several “read-outs” which track non-stationary reward environments, with the option for arbitration based upon the precision of each neural circuits estimate.

In the study of Humphries et. al., (2012) varying the level of tonic dopamine influenced explore-exploit decisions in classical two choice, fixed contingency, probabilistic tasks (Frank, 2005). In their simulations, a range of dopamine levels were tested, but remained at constant non-fluctuating values between trials. The closest equivalent model in this study failed to track the high value choice in the restless bandit task at better than chance levels. This result emphasises the additional computational value of tonic dopamine fluctuations on a fine time scale. Several experimental findings would support that this is indeed the case *in vivo*. Voltametry recordings of extracellular dopamine release in the Nucleus Accumbens (NAc) exhibit fluctuations at subs-second temporal resolution that exceeds the trial-to-trial resolution required for our model (Berke, 2018; Mohebi *et al*., 2019). Tonic dopamine efflux also increases in the NAc when the reward schedule is more unpredictable and reduces to baseline levels when the reward payout regularises (St Onge *et al*., 2012). These higher levels of striatal tonic dopamine are also paralleled by increased tonic firing rates in single VTA neurons when reward delivery is unpredictable (Fiorillo *et al*., 2003; de Lafuente & Romo, 2011). In humans, striatal activity is also greatest when the reward schedule unpredictable (Preuschoff *et al*., 2006).

In their account of tonic dopamine, (Niv *et al*., 2007) proposed this tracked the average reward rate, with higher levels of tonic dopamine leading to increased rate of responding rates and response vigour. At first glance, our proposal of high tonic dopamine during exploration, where reward payout is lowest, would seem to contradict this idea. A closer consideration of their theory reveals that this is applicable to free-operant tasks, making direct comparison to a fixed response, trial based schedule, such as the restless bandit task more difficult. We would also argue that there is a strong biological incentive to increase response rate and response vigour during explorative decisions, to rapidly eliminate low value and efficiently identity the highest value choice to exploit. Equally, during exploitative decision making, there seems little point, and greater cost, to invigorating the rate of responses when the availability of rewards is predictable. A basic prediction that would reconcile both our and Niv’s theory would be the increased response rates during higher uncertainty, in a free operant task with a high payout volatility. The finding of reduced reaction times during explorative decisions would suggest that uncertainty is associated with increased vigour of responding (Gershman, 2019).

### Relationship to experimental data

The central idea of our model is that increasing the background level of dopamine raises the stochasticity of action selection (random exploration) as the basal ganglia’s ability to select deterministically is reduced. “Soft” selection was proposed by Suryanarayana et al. (2019) as a potential source of exploration, as higher levels of dopamine reduces basal ganglia’s capacity to filter cortical input leading to a greater number of actions being disinhibited. Animals genetically engineered to be hyper-dopaminergic, exhibit behaviour that is consistent with increased random exploration (Zhuang *et al*., 2001), however, this behaviour could result from influence from extra-striatal sites such as prefrontal cortex (Beeler *et al*., 2010). Increased directed exploration, towards novel stimuli, has been also observed animals with increased striatal tonic dopamine. This effect may not directly compared with the random exploration seen in our model as distinct neural mechanisms may contribute to these different forms exploratory behaviour (Costa *et al*., 2014; Wilson *et al*., 2014). Pharmacological studies in humans which increase extracellular levels of dopamine also increase explorative behaviour, particularly in genetically susceptible individuals (Kayser *et al*., 2015; Gershman & Tzovaras, 2018). Consistent with our predictions, the increase in exploration in a foraging task, caused by administering a dopamine receptor agonist, was most marked when foraging for in an environment where reward was more uncertain (Le Heron *et al*., 2020). Paradoxically, in the context of these findings, administration of dopamine receptor antagonists has also been linked to increased exploration. In their analysis, using a combination of experimental and mathematical approaches, Cinotti et. al., (Cinotti *et al*., 2019) demonstrated that this effect was best explained by reduced the amplitude of phasic reward prediction error signalling. A similar blunting of phasic RPE mediated action-value mapping could also explain increased random exploration observed in animals with genetically impaired D1 and D2 dopamine signalling (Kwak *et al*., 2014; Cieslak *et al*., 2018). The effect of genetic factors which are associated with increased exploitative learning is consistent with their influence on augmenting phasic RPE signalling (Frank *et al*., 2009b), but is harder to reconcile with data suggesting greater stochasticity of choice with increased D2R binding in PET studies (Adams *et al*., 2020). The results of Chakroun et. al (Chakroun *et al*., 2020), who found no effect of dopamine antagonism or Levodopa on random exploration, but a reduction in directed exploration on Levodopa, also illustrate how difficult it is to experimentally separate, and accordingly interpret, the differential effects of phasic and tonic dopamine signalling on exploratory behaviour. Paradoxically therefore, reducing phasic RPE signalling, or increasing background tonic dopamine may lead to the same behaviour, an increase in random exploration. Whether this is by reducing the selectivity of the basal ganglia output, as proposed here, or by “washing out” fine phasic signalling due to saturation (Beeler *et al*., 2010) or a mixture of both, is unclear, but either mechanism may contribute to the inverted U-shaped relationship between dopamine and learning (Clatworthy *et al*., 2009). The best performing model reproduced this inverted U-shaped relationship (Figure 5A & C), emphasising that a relatively narrow range of background changes in dopamine level are required for optimal explore-exploit decisions. If this range is either non - existent (for example our “fixed” model with no dopamine fluctuations) or too broad, with an upper limit of dopamine that is excessive, learning and performance degrades accordingly.

In agreement with other proposals (Friston *et al*., 2012), our model would also predict an additional influence of tonic dopamine at the level of how the RPE signal influences cortico-striatal synaptic gain. The neuromodulatory effects of dopamine on cortico-striatal plasticity in our model are governed by the dopamine weight change curve (Gurney *et al*., 2015) which assumes dopamine promotes synaptic potentiation at D1 and depotentiation at D2 receptors (Shen *et al*., 2008). Here we found that the models explore-exploit performance was enhanced when the RPE’s influence on plasticity was interpreted relative to the background level of dopamine. This mechanism assumes that the gain of these two signals can be regulated independently of one another (Grace *et al*., 2007; Schultz, 2007), with the effect of making tonic dopamine analogous to a background contrast to the more temporally refined phasic RPE signal. Accordingly, higher background levels of dopamine enhanced synaptic potentiation in D2R, as these were more sensitive to excitatory phasic pauses, and less sensitive to inhibitory bursts, when under greater tonic inhibitory dopaminergic tone. The opposite relationship was found when dopamine levels were at the lower bounds of the optimum range. The effect of this interaction would be to increase in indirect pathways activity during exploratory/soft action selection and in turn prevent repetition of low value choices (Kravitz *et al*., 2012). It would also be consistent with the experimental effects of optogenetic stimulation of indirect pathway neurons on promoting switching behaviour (Nonomura *et al*., 2018). At the optimal lower bounds of tonic dopamine, exploitative behaviour was mediated by combination of hard selection (increased selectivity) and increased gain at direct pathway cortico-striatal synapses. This increased direct pathway excitability during exploitation is supported by the experimental findings of enchanced reinforcement learning and choice perseveration when this pathways excitability is increased (Kravitz *et al*., 2012; Nonomura *et al*., 2018).

A significant assumption of this work is that the basal ganglia can access information on a trial-to-trial basis about the reward uncertainty. Uncertainty representation in the striatum has recently been supported by the modelling studies (Mikhael & Bogacz, 2016) and experimentally by the finding of a group of striatal neurons which encode this in their firing rates (White & Monosov, 2016). Equally, the basal ganglia may receive this information from neurons which have been shown to encode uncertainty in the ACC (Behrens *et al*., 2007; Rushworth & Behrens, 2008) or dorsolateral prefrontal cortex (Tomov *et al*., 2020). Where or how the uncertainty estimate is generated, may be less important to the predictions of our model than that the basal ganglia have access to this information. For this to influence the level of tonic dopamine, the basal ganglia’s uncertainty estimate must also be capable of self-regulating tonic dopamine release, presumably by influencing the excitability of the VTA. The most likely candidate for this would be the inhibitory influence of the Ventral pallidum (VP) on the VTA which was proposed to provide an independent source of controlling tonic dopamine levels (Floresco *et al*., 2003). The baseline firing rate of VTA neurons during the “uncertainty response” (Schultz, 2007; Schultz *et al*., 2008) increases two fold and has been predicted to increased extracellular striatal dopamine by ~10–30 nM. This would be expected to preferentially inhibit the indirect pathway and lead to necessary softening of the action selection which we predict is one source of random exploratory decision “noise.”

A limitation to our model is the omission of any details pertaining to neuromodulators such as noradrenaline and acetylcholine in uncertainty estimation (Yu & Dayan, 2005). Of particular relevance is cholinergic feedback circuit formed by cholinergic Tonically Active Neurons (TANs) (Franklin & Frank, 2015). Due to our focus on the action selection function of basal ganglia, we necessarily omitted the biophysical detail required to explore their detailed mechanisms of uncertainty estimation. Future work may allow us to synthesise both these local striatal cholinergic and extra-striatal dopaminergic feedback circuits, which may exert distinct influences on cortico-striatal plasticity and selectivity of the basal ganglia circuit as a whole. We also did not include any detailed description of how tonic dopamine levels ramp up or decay between trials, adopting a simplified binary state. Our assumption is that in fixed stimulus-response, trial-based tasks, how background dopamine changes between the stimulus and the response is irrelevant providing that it reaches the optimal level by the point of action selection. In tasks which rely upon free, internally generated responses, this assumption is much less likely to be justified and more physiologically plausible dopamine fluctuations (such as ramping) are likely to be necessary and may be computationally valuable.

### Experimental predictions

Recent experimental data has questioned the validity of separate and independently controllable phasic and tonic dopamine systems for learning. In part, this has been due to the absence of any clear evidence of behavioural correlate of changes in tonic VTA firing (Mohebi *et al*., 2019). One explanation for this is that slow trial-to-trial fluctuations in the baseline firing rates (and corresponding levels of striatal tonic dopamine) only become relevant to learning under conditions when the computational burden and corresponding volatility of the reward schedule are sufficiently high. This would explain the lack of clear behavioural effects of manipulations that would be expected to modify tonic dopamine when classical two-choice, fixed contingency tasks are studied (Vancraeyenest *et al*., 2020). When the volatility of the reward schedule becomes close to those seen here, or in natural environments, such as foraging (Le Heron *et al*., 2020), a circuit which tracks the uncertainty of the reward and feeds this back to control the background firing rate of the VTA, would be a source of random exploratory decision “noise,” by influencing the selectivity of action selection in the basal ganglia. Modulation of the excitability of afferent (e.g VP) or efferent (e.g. indirect pathway MSNs) limbs within this circuit could causally test our hypothesis in the appropriate behavioural context in animals. Our model might also offer a further explanation as to why loss of dopamine associated with Parkinson’s disease (PD) leads to apathy (Sinha *et al*., 2013; Husain & Roiser, 2018) and why this can be improved with dopamine agonists (Thobois *et al*., 2013). Dopamine depletion in PD would impair fluctuations in tonic dopamine that are necessary in our model to invigorate random exploration. This prediction requires relies upon further understanding as to the extent to which apathy and impulsivity can be explained by shared neurobiological mechanisms that underlie explore/exploit decisions (Addicott *et al*., 2017). This relationship is unknown in PD but would be supported by the finding of reduced exploration correlating with levels of apathy in other patient groups (Batrancourt *et al*., 2019).

### Conclusions

The exact computational function of tonic dopamine is unclear. Here we demonstrate that explore exploit decisions in a non-stationary, reward environment characterised by high pay-out uncertainty, can be optimally by the basal ganglia’s action selection circuit. This circuit’s performance is dependent upon the interaction between tonic levels of dopamine on the selectivity of the action selection circuit and gain of striatal synaptic plasticity.

## Abbreviations

MSN: Medium Spiny Neuron
RPE: Reward Prediction Error
GPe: Globus Pallidus externa
GPi: Globus Pallidus interna
D1R: Dopamine receptor type 1
D2R: Dopamine receptor type 2
LTP: Long Term Potentiation
LTD: Long Term Depression

## Acknowledgements

We thank Mark Humphries and Rafal Bogacz for informative discussions on explore-exploit decision making and feedback on earlier version of this manuscript.

## Appendix

### Activation and output functions for basal ganglia action selection circuit (GPRe)

Here we summarise the basic output and activation functions for the full model described in detail in Suryanarayana et. al., (2019). All values of the constants for these are included in Appendix table 1. From striatal activation function in the main methods section (Equation 1), the striatal activity in channel *i* for the general case *n* is represented by 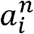 The output relationship for the striatal neurons is then;

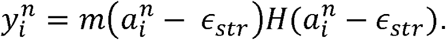

The activation function for the STN is;

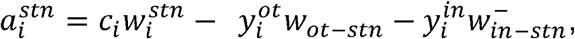

And its output relationship is 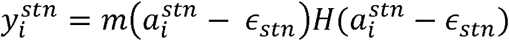.

As per the model schema (appendix figure 1B) the extended model includes the different neural populations of the globus pallidus externa (GPe), including the “outer”, “inner” subdivision of the prototypical GPe cells (GP-TI) which project to the GPi as well as the striatum and the arkypallidal cells (GP-TA) which project back to the striatum. The GPe outer neurons activation function is;

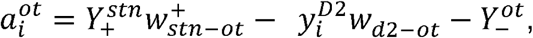

Here, 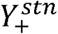 represents the diffuse excitatory input from the STN where 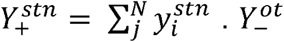 is the effect inhibitory intrinsic collaterals, where 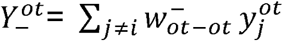 and 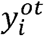 the GPe outer neurons output function is: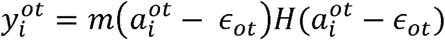.

The GPe inner neurons activation function is;

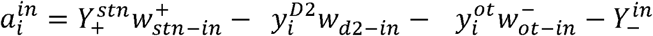, with 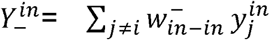. The output function is then 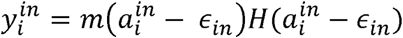.

Finally, the GPe-TA cells activation is;

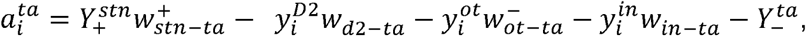

Where 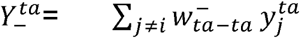 and the corresponding output function is 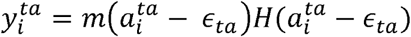.

### Activation and output functions for basal ganglia action selection circuit (GPR)

As per the original GPR model (Gurney et. al., 2001) the striatal activity is 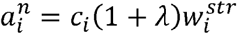. Again *λ* here represents the tonic level of dopamine and takes on positive values for the direct pathway and negative values for the indirect pathways striatal activation.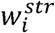 represents the cortico-striatal weights which all start at values of 1 for both direct and indirect pathways but vary according to the dopamine-weight change curve (equation 5). Output functions *y*_i_ are calculated using the same general form as for the GPRe model. The STN activity is defined as:

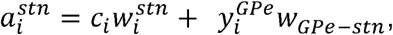

where *w*_*GPe_ stn*_ = -1, 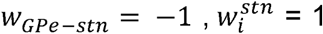, and 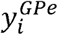 is the output function of the GPe. The activity of the GPe is;

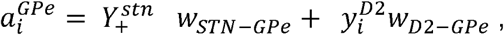

Here *w*_*STN_GPi*_,*w*_*D2-Gpe*_ are set at values of 0.9 and -1,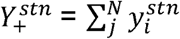 Finally, the GPi’s output is defined as ;

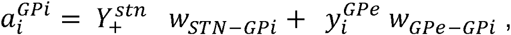

The values for *w*_*STN_Gpi*_, and *w*_*GPe-Gpi*_ are 0.9 and -0.3 respectively.

**Appendix Table 1.**

| Synaptic weights | Values |
| --- | --- |
| $w_{D2-ot}$ | -1 |
| $w_{D2-in}$ | -1 |
| $w_{D2-ta}$ | -1 |
| $w_{D1-GPi}$ | -1 |
| $w_{ot-D1}$ | 0.5 |
| $w_{ot-D2}$ | 0.5 |
| $w_{in-D1}$ | 0.25 |
| $w_{in-D2}$ | 0.25 |
| $w_i^{stn}$ | 1 |
| $w_{stn-GPi}$ | 0.9 |
| $w_{stn-ta}$ | 0.8 |
| $w_{ot-stn}$ | -0.8 |
| $w_{ot-GPi}$ | -1 |
| $w_{stn-ta}$ | 0.8 |
| $w_{ot-ot}^-$ | -0.75 |
| $w_{in-in}^-$ | -0.75 |
| $w_{ta-ta}^-$ | -0.75 |
| $w_{ot-in}^-$ | -0.3 |
| $w_{ot-ta}$ | -0.75 |
| $w_{in-ta}$ | -0.75 |
| $w_{ta-D1}$ | -0.25 |
| $w_{ta-D2}$ | -0.25 |

**Appendix Table 2.**
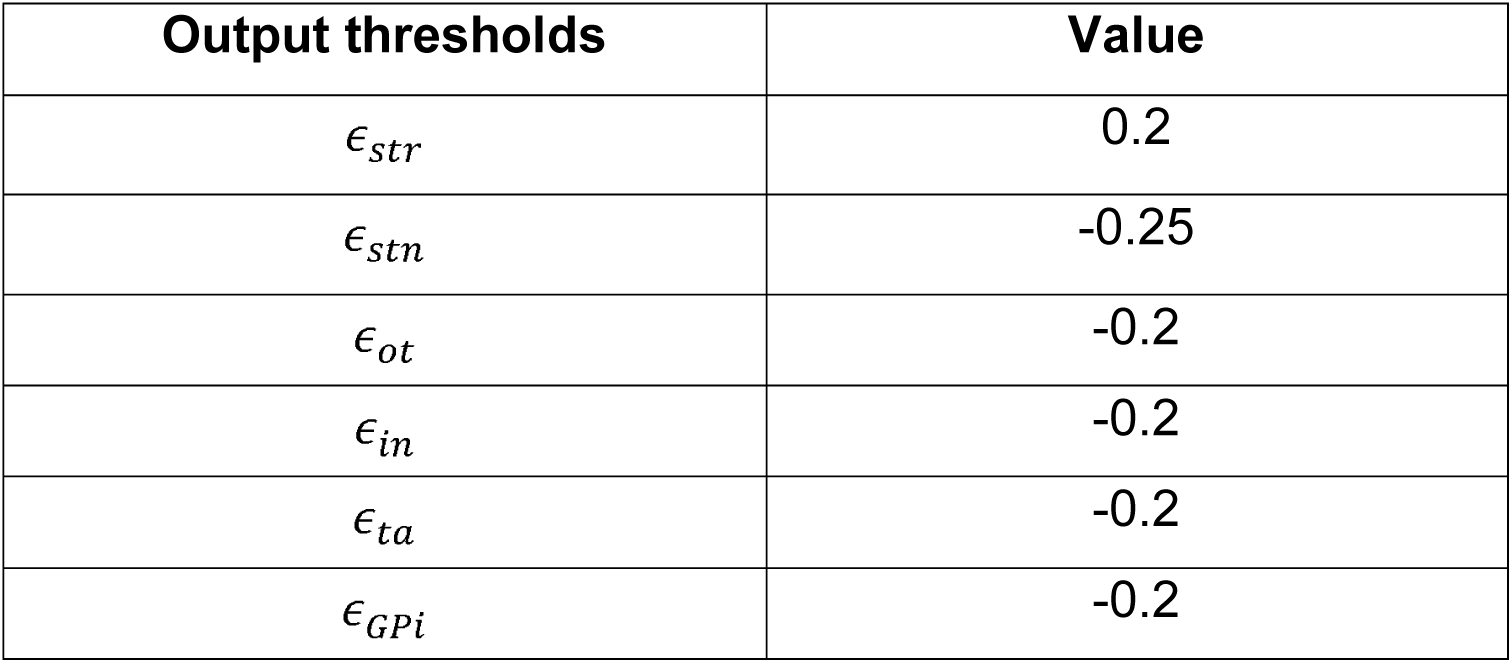

**Figure 1 Appendix.**
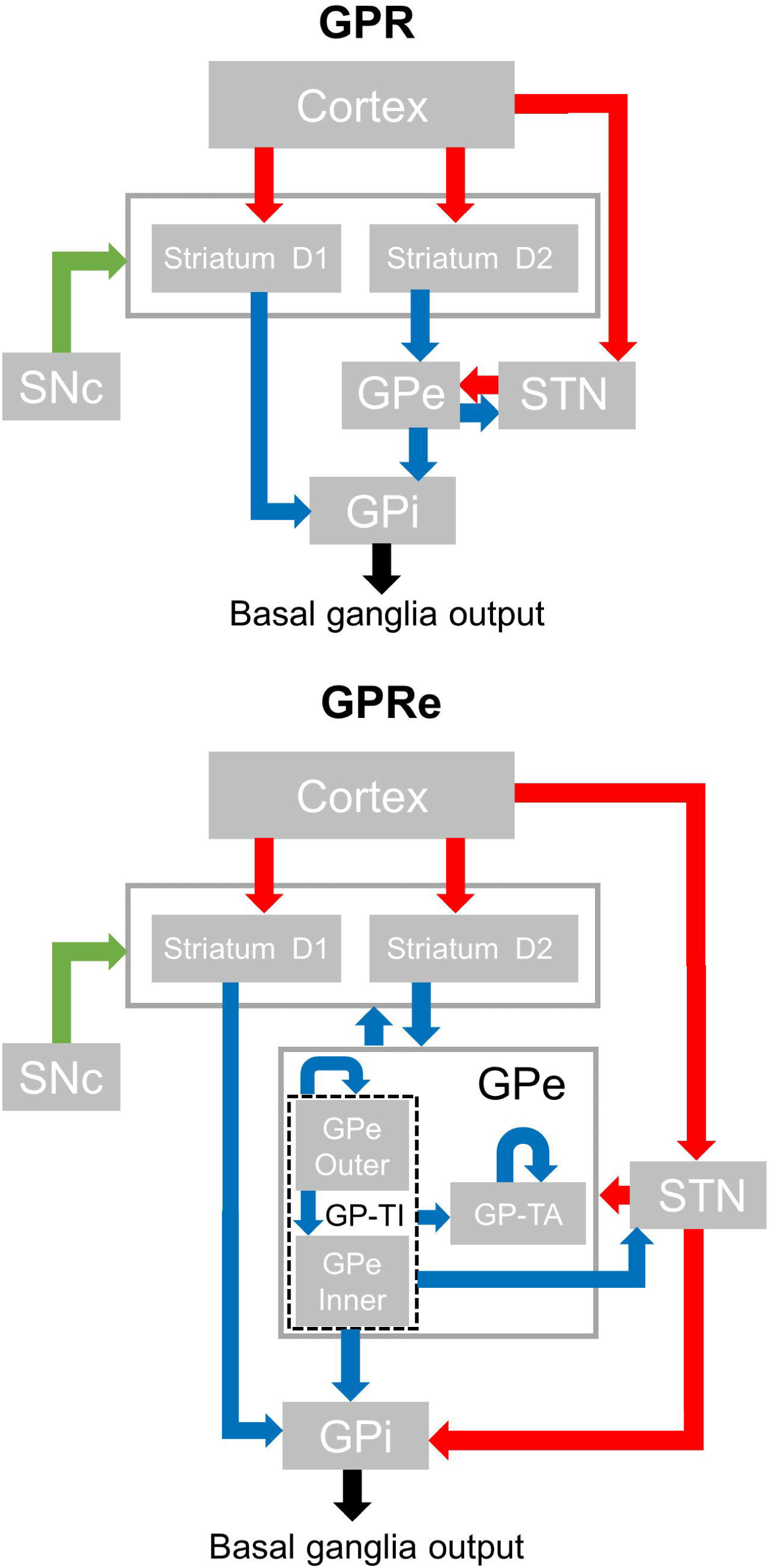
Model schema for the basal ganglia’s action selection circuit for the Gurney-Prescott-Regrave model (GPR) in **(A)** and the extended (GPRe) version of this model which includes updated intrinsic GPe connectivity (Suryanarayana et. al., 2019) in **(B)**. This includes partitioning of the GPe into both inner (GPe Inner) and outer (GPe Outer) populations and designation of Arkypallidal (GP-TA) and Prototypical GPe (GP-TI) subpopulations. Green arrows represent dopaminergic neuromodulatory projections, red glutamatergic excitatory and blue gabaergic inhibitory projections. GPi: globus pallidus interna, GPe; Globus pallidus externa, STN, Subthalamic nucleus.

